# Obstructive Sleep Apnea Improves with Non-invasive Hypoglossal Nerve Stimulation using Temporal Interference

**DOI:** 10.1101/2023.04.06.535917

**Authors:** Florian Missey, Malin Silverå Ejneby, Ibrahima Ngom, Mary J. Donahue, Jan Trajlinek, Emma Acerbo, Boris Botzanowski, Antonino M. Cassarà, Esra Neufeld, Eric D. Glowacki, Lee Shangold, William M. Hanes, Adam Williamson

**Affiliations:** International Clinical Research Center, St. Anne’s University Hospital Brno, 60200 Brno, Czech Republic; Institute de Neurosciences des Systèmes (INS), INSERM, Aix-Marseille Université, 13005 Marseille, France; Department of Biomedical Engineering, Linköping University, 58185 Linköping, Sweden; Laboratory of Organic Electronics, Campus Norrköping, Linköping University, 602 21 Norrköping, Sweden; Central European Institute of Technology, Brno University of Technology, 61200 Brno, Czech Republic; IT’IS Foundation for Research on Information Technologies in Society, 8004 Zurich, Switzerland; ENT and Allergy Associates, 1500 Route 112, Port Jefferson Station, NY 11776, USA; Somnial Inc., 25 Health Sciences Dr, Stony Brook, NY, USA

**Keywords:** Temporal Interference, peripheral nerve stimulation, Hypoglossal Nerve, Obstructive Sleep Apnea

## Abstract

**Background:** Peripheral nerve stimulation is used in both clinical and fundamental research for therapy and exploration. At present, non-invasive peripheral nerve stimulation still lacks the penetration depth to reach deep nerve targets and the stimulation focality to offer selectivity. It is therefore rarely employed as the primary selected nerve stimulation method. We have previously demonstrated that a new stimulation technique, temporal interference stimulation, can overcome depth and focality issues.

**Methods:** Here, we implement a novel form of temporal interference, bilateral temporal interference stimulation, for bilateral hypoglossal nerve stimulation in rodents and humans. Pairs of electrodes are placed alongside both hypoglossal nerves to stimulate them synchronously and thus decrease the stimulation amplitude required to activate hypoglossal-nerve controlled tongue movement.

**Results:** Comparing bilateral temporal interference stimulation with unilateral temporal interference stimulation, we show that it can elicit the same behavioral and electrophysiological responses at a reduced stimulation amplitude. Traditional transcutaneous stimulation evokes no response with equivalent amplitudes of stimulation.

**Conclusions:** During first in-man studies, temporal interference stimulation was found to be well-tolerated, and to clinically reduce apnea-hypopnea events in a subgroup of female patients with obstructive sleep apnea. These results suggest a high clinical potential for the use of temporal interference in the treatment of obstructive sleep apnea and other diseases as a safe, effective, and patient-friendly approach.

**Trial registration:** The protocol was conducted with the agreement of the International Conference on Harmonisation Good Clinical Practice (ICH GCP), applicable United States Code of Federal Regulations (CFR) and followed the approved BRANY IRB File # 22-02-636-1279.

## BACKGROUND

Obstructive sleep apnea (OSA) is a highly prevalent disorder characterized by episodes of decreased or absent inspiratory airflow during sleep^29,1^. This can lead to excessive daytime sleepiness and other comorbidities such as stroke, hypertension, or depression. To prevent airway collapse or narrowing during sleep, positive airway pressure delivered via a well-fitting mask is used as the primary treatment for patients with moderate to severe OSA^8,24^. However, adherence to the therapy remains limited despite advances in quieter pumps or softer masks^24^. For this reason, hypoglossal nerve stimulation (HNS), which modulates the airway through neural stimulation of the genioglossus muscle (causing tongue protrusion), has been developed as an alternative treatment^25,23,27,30,9^. To date, HNS is an invasive technique that requires an electrode in direct contact with the hypoglossal (cranial XII) nerve, as well as a battery placed in the chest to run the electrical stimulation^19^. Avoiding surgeries would be the best way to prevent adverse effects from the surgery itself, but also those related to the electrodes placed inside the body^3^. Patient exclusion, anesthesia, the manipulation of the nerve, and the rejection of the electrode/battery inserted in the patients could potentially be overcome by using non-invasive techniques^4^.

Direct nerve stimulation using topical electrodes placed on the skin is possible – currents can reach the nerve at depth without the need for invasive surgery^26,18,13^. However, the use of Transcutaneous Electrical Nerve Stimulation (TENS) is not very efficient and requires a high current amplitude to reach the nerve target. In most cases, this leads to an unpleasant feeling because of the high current density applied on the skin, which again could reduce adherence to therapy. Non-invasive temporal interference (TI) stimulation can help overcome this issue by enabling depolarization of deep nerves with modulated electrical fields without activation of overlaying structures. Even though these structures are exposed to higher field strengths, electrophysiologicaly active tissues do not efficiently respond to kHz frequencies; however, they can respond to the low-frequency modulation generated by TI, which is able to selectively focus to activate the target nerves) It uses two high frequency waves that mix at the targeted location, for example, a brain region or the location of a peripheral nerve.. TI stimulation has previously been shown to be useful for brain stimulation^12,22^. Recently we have shown that temporal interference can be applied with topical electrodes placed on the skin to stimulate the sciatic nerve of mice without eliciting contraction of non-targeted muscles or skin pain and damage^6^.

In the present work, we show that the use of cutaneous electrodes can lead to an efficient stimulation of the hypoglossal nerves in mice using temporal interference stimulation. We used two high-frequency carriers, at *f1* and *f2 = f1 + Δf* Hz to create an *f* Hz stimulation at the location of the hypoglossal nerve. The hypoglossal nerves are nonsuperficial, located beneath layers of muscle, making them difficult to selectively target with single-channel transcutaneous electrical stimulation. To realize a highly selective hypoglossal nerve stimulation, we provide bilateral TI (bTI) which allowed us to decrease the exposure amplitude on each side and thus minimize stimulation of neighboring tissues while eliciting strong nerve stimulation-induced muscle movement. In addition to decreasing the amplitude of each stimulation pair, using bTI for hypoglossal nerve stimulation induces a central tongue protrusion (compared to lateral stimulation that induces lateral protrusion^9,15^ and optimizes airflow. Moreover, controlling field orientation is also crucial in peripheral nerve stimulation. Aligning temporal interfering fields alongside the nerve enhances their nerve depolarization efficiency and helps nerve recruitment^6^. Finally, using a highly precise temporal interference protocol with cutaneous electrodes placed along the nerve, we were able to provide safe and completely tunable hypoglossal nerve stimulation for OSA patients, with a concomitant reduction of symptoms in a subset (**Figure 1**). Here we highlight the use of bTI for obstructive sleep apnea treatment, but temporal interference can be used in various protocols to provide on-demand stimulation for peripheral nerves without a need for device implantation.

**Figure 1.**
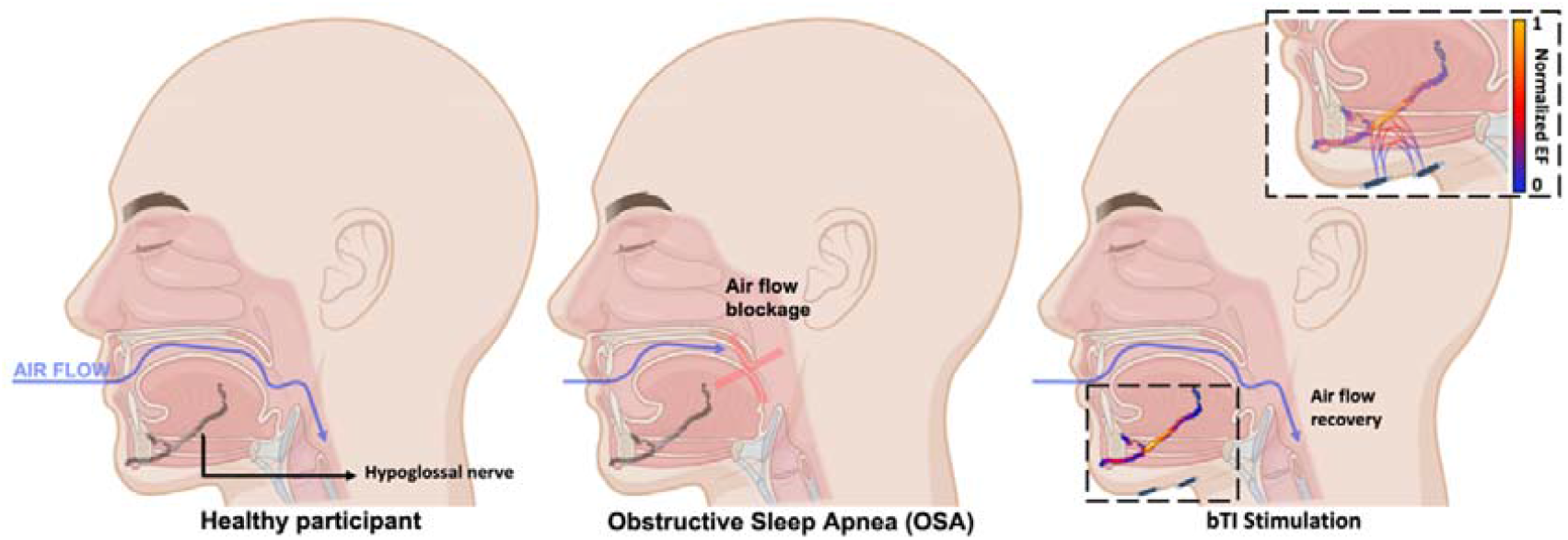
Airflow recovery in obstructive sleep apnea patients using bilateral temporal interference stimulation. In healthy people, air flows normally during sleep with no obstruction of the airway between the mouth and the lungs. Patients suffering from sleep apnea have a blockage of the upper airway due to the collapse of the tongue in the back of the throat during sleep, preventing patients from breathing normally. Using bilateral temporal interference stimulation of the hypoglossal nerves, which controls tongue tone and movement, prevents it from collapsing and allows recovery of airflow. The peak-normalized electric field presented in the inset was calculated using finite element modeling (FEM) electro-quasistatic modeling (Sim4Life, ZMT Zurich MedTech AG, Switzerland).

## METHODS

### Animal study

#### Animals

Experiments were realized using a mouse model under the agreement of European Council Directive EU2010/63 and French Ethics approval (comité d’éthique en experimentation animale n°70 Williamson, n. APAFIS 20359 - 2019041816357133). 8x OF1 mice, 8 to 10-week-old, (Oncins France 1, Charles Rivers Laboratories, France) underwent the stimulation protocol presented in this study. Mice were housed in cages of 4 and under a normal 12/12h day-night cycle at room temperature and with water and food *ad libitum*.

#### Animals’ preparation

Mice received an intraperitoneal injection (2.5µL/g) of 1% xylazine (20mg/kg) and 99% ketamine (50mg/kg) mix. The mice’s temperature was monitored at every step of the experiment, and we took care of covering the mice’s eyes with vitamin A to avoid damage. Mice were placed on their back to have an accessible stimulation window around both hypoglossal nerves. Once placed in a comfortable position, the throat was depilated using hair removal Veet® cream. We let the cream stand for five minutes maximum to avoid skin inflammation and removed it gently using water.

#### Electrical stimulation using golden stimulation electrodes in mice

Once mice preparation was complete, 4 gold-plated pin headers (Farnel® ref. 825433-4) were placed on one side of the throat, above the location of the hypoglossal nerve. Once the position of the 4-golden pin apparatus was well aligned to the hypoglossal nerve location, the apparatus was gently pushed toward the mouse skin until all 4 pins were touching it. Temporal Interference stimulation was provided using two waveform generators (Keysight EDU33212A from Keysight®) connected to voltage to current converters (DS5 isolated bipolar constant current stimulation from Digitimer®). Each TI stimulation pair (composed of a stimulation pin and a ground pin) was connected to its function generator and DS5. A standard TI stimulation with a 5Hz offset (3000 and 3005Hz) and 1mA per pair was realized to evoke a tongue movement. If no behavioral output was visible, electrode placement was slightly moved until the stimulation elicits a 5Hz tongue movement. Finally, another stimulation apparatus was placed on the other side of the throat to target the second hypoglossal nerve and provide a bilateral TI stimulation.

#### CMAP recordings in mice

A thin 70µm platinum wire was insulated, except for the tip, and this tip was inserted in the tongue muscle to act as recording electrode with another wire placed as a reference in the *tibialis anterior* muscle. Recordings were acquired thanks to an RHS 128Ch Stim/Recording System (Intan technologies®) with an internal 50Hz notch filter to cancel ambient noise

*Force recordings in mice:* Efficient bTI stimulation of the HN is characterized by tongue twitching at the offset frequency of the bTI stimulation. To record the tongue twitching and its force we used an FT50 isometric force transducer (maximal load ± 50cN) coupled to an HMO3004 Series Oscilloscope (ROHDE & SCHWARZ®).

#### Statistical analysis

Both CMAP recordings and force recordings were plotted and analyzed using MATLAB 2021 (MathWorks). All CMAP recordings were pre-filtered with a bandpass filter (1-4000Hz) to reduce noises and frequency plots were generated (spectrogram and periodogram) to highlight the global frequency of tongue CMAPs. Once the global frequency has been identified (around 200Hz), all recordings were filtered using another bandpass filter (100-400Hz) for better visualization of the individual CMAPs. To characterize tongue CMAPs, three intrinsic components were analyzed over 30 CMAPs per group: Intrinsic Frequency, Duration, and Amplitude. These variables were extracted and compared using first normality tests (Shapiro) for each group and then non parametric tests (Wilcoxon-Mann-Whitney and Friedman, power = 0.8) to compare the different groups (unilateral TI versus bilateral TI)

### Early feasibility study in OSA patients

#### Participants

To be enrolled in this study, patients had to be 18 years old or above, in general good health, and diagnosed with moderate or severe sleep apnea based on an in-sleep laboratory polysomnogram (PSG) recorded not longer than 6 months ago in a United Sleep Diagnostics (Commack, NY) facility and documenting an AHI of 15 to 50/hour, and willing to abstain from alcohol and caffeine on the day of the study. A total of 12 patients were included in this early feasibility study (7 men and 5 women), aged 48.1 ± 10.7 years and with an average BMI of 30.6 ± 6.8kg/m^2^ (**Table 1**).

**Table 1.** Participant characteristics.

| Characteristic | Participants (n=12) |
| --- | --- |
| Age | 48.1 ± 10.7 yrs |
| BMI* | 30.6 ± 6.8 kg/m<sup>2</sup> |
| Gender – Male | 7 (58%) |
| Gender – Female | 5 (42%) |
| Baseline AHI | 30.1 ± 11.3/hour |
| *The body-mass index is the weight in kilograms divided by the square of the height in meters. | means ± SD |

#### Exclusion criteria

Patients were excluded from this early feasibility study if they were diagnosed with no, mild, or too severe OSA (Apnea Hypopnea Index > 50). In addition, patients with a Body Mass Index under 18.5 kg/m^2^ and over 32 kg/m^2^ were excluded from the present study, due to their decreased probability of responding to therapy. Finally, patients were excluded if pregnant, or when having one or more of the following diseases: enlarged tonsils (size 3-4) and/or adenoids, nasal polyps, neuromuscular diseases, hypoglossal nerve palsy, abnormal pulmonary function, severe pulmonary hypertension, valvular heart disease, heart failure (New York Heart Association, NYHA III IV), recent myocardial infarction, significant cardiac arrhythmias, atrial fibrillation, stroke, opioid usage, uncontrolled hypertension, known allergic reaction to adhesives or latex, active psychiatric disease, co-existing non-respiratory sleep disorder, or significant metal implants including pacemakers.

#### Study design

This early feasibility study was designed as an open-label, monocentric, single-group treatment trial where patients were under their own control. The protocol was conducted with the agreement of the International Conference on Harmonisation Good Clinical Practice (ICH GCP), applicable United States Code of Federal Regulations (CFR) and followed the approved BRANY IRB File # 22-02-636-1279.

Qualified patients underwent a night PSG wearing the stimulation device for bilateral hypoglossal nerve stimulation. The device consists of 4 pairs of commercially available gel-based electrodes, 2 pairs for each HN placed on the corners of a 2cm square around the digastric anterior belly muscle and oriented to be aligned with the HN. The 8 electrodes in total were placed at the bedside by the technician, once the patient was ready to go to bed. Stimulation was applied through the pairs of electrodes, as described in Figure 4. Before going to sleep, the patients were subjected to an amplitude titration (from 0.1 to 8mA) to evaluate what stimulation amplitude was needed to elicit a tongue movement; when the movement threshold was reached, the stimulation was turned off and the patient could go to sleep. Once the patient was asleep, the stimulation was switched on at the lowest amplitude (1mA) and ramped up by steps of 0.5mA until the pre-defined threshold (patient-specific) was attained. Any time during the night, the patient could ask to pause the treatment and be disconnected from the stimulators, to use the bathroom for example. Lastly, the patients were given a questionnaire at the end of the night to relate their experience with the stimulation device.

#### Outcomes measures

The primary outcome of the study was to evaluate the efficiency of the stimulation device through the measurement of AHI during the in-laboratory investigation. A positive endpoint was defined as a decrease of the AHI of at least 50% with an overnight AHI of 20 or less. The secondary outcomes included Percentage Sleep Time at SaO2 < 90% (determined by the time below a SaO2 level of 90% compared to that at baseline) and Safety based on a description of all reported adverse events. The tertiary outcome focused on exploring patients’ comfort when wearing the stimulation device overnight using retrospective questionaries.

#### Statistical analysis

For the primary outcome, sleep summaries were analyzed in a single-blinded approach under Rule 1A of the AAAS guidelines (apneas are identified as ≥ 90% flow reduction events lasting more than 10 seconds; hypopneas are defined as a ≥ 30% flow reduction for more than 10 seconds with a decrease of 0_2_ saturation ≥ 3%), and the sleep technician who analyzed the sleep summaries was not aware of the stimulation status. The obtained AHIs were compared to the ones obtained during the baseline (the initial PSG established within 6 months before the present study). To determine whether the therapy was efficient, Wilcoxon tests were used (if the compared groups were not normally distributed, i.e., p-value_Shapiro test_ < 5%; ⟨ = 1%) or by Student’s t-test (if the compared groups were normally distributed, i.e., p-value_Shapiro test_ > 5%; two-tailed, paired).

### In silico simulations

#### Finite element simulations

Simulations of temporal interference were performed on the Sim4Life software (Zurich MedTech AG), using the electro-ohmic quasi-static solver, which solves the equation ∇σ∇⍰ = 0, where σ is the local electrical conductivity, ω the angular frequency, and the E field is obtained as E=-∇⍰. This approximation of Maxwell’s equations is suitable because σ≫ω⍰ (ohmic currents dominate displacement currents) and the wavelength is much longer than the exposed domain, and it is numerically superior when the discretization is much finer than the wavelength. The E-field is only determined in the lossy domain (σ≠0) – thus, the default grid covers only the lossy domain. The mouse and human models are part of the Virtual Population (Male PIM1 Mouse for the mouse and Jeduk V4.0 for the human model) and the simulation platform automaticaly assigns tissue properties according to the IT’IS Foundation database of tissue properties^14^. The mouse hypoglossal nerve was modelled by extruding a circular cross section (diameter: 0.2mm) along the trajectories in the model and assigning anisotropic conductivity tensors (longitudinal and transversal conductivity; main conductivity axis aligned with the gradient of an auxiliary diffusion simulation along the nerve geometry). Electrodes were modeled as perfect electrical conductors (PEC) and their placement was varied to identify optimized exposure conditions for hypoglossal nerve targeting. The stimulation was performed with Dirichlet boundary conditions at active electrodes and the results were total-current-normalized. The maximum envelope modulation amplitude was computed using the formula from **Grossman & al**., **2017** (Equation 1).

## RESULTS

### Bilateral TI can evoke tongue activities with less current than regular TI in mice

It is well known that stimulation of one, or two, hypoglossal nerve(s) generates tongue movement. Using this tongue movement as a direct and positive behavioral output for HN stimulation, we used two techniques to monitor the impact of our temporal interference stimulation protocols: electromyography of the tongue muscle (EMG) and force recording of the tongue movement (FR). Before any stimulation, a baseline recording of EMG and FR was performed. No tongue movements were visible, and we did not observe any muscle activity in the EMG recording – only breathing artifacts were visible. To stimulate hypoglossal nerves and elicit tongue movement, temporal interference was administrated using a bilateral TI apparatus. Each apparatus was composed of two pairs of electrodes, aligned along the respective hypoglossal nerves and providing 3000 (*f1)* and 3005Hz (*f2 = f1+Δf, Δf* = 5Hz) exposure respectively, to generate a 5Hz field targeting the HN (**Figure 2**). For regular TI stimulation, only one apparatus was active (i.e., only one HN was stimulated, **Figure 2B-C**) whereas, in the bTI protocols, both apparatuses were active (i.e., both HN were stimulated, **Figure 2D**). In all cases, stimulation amplitudes were increased in steps of 100µA until a clear tongue movement could be recorded (tongue movement of around 20cN on the FR).

**Figure 2.**
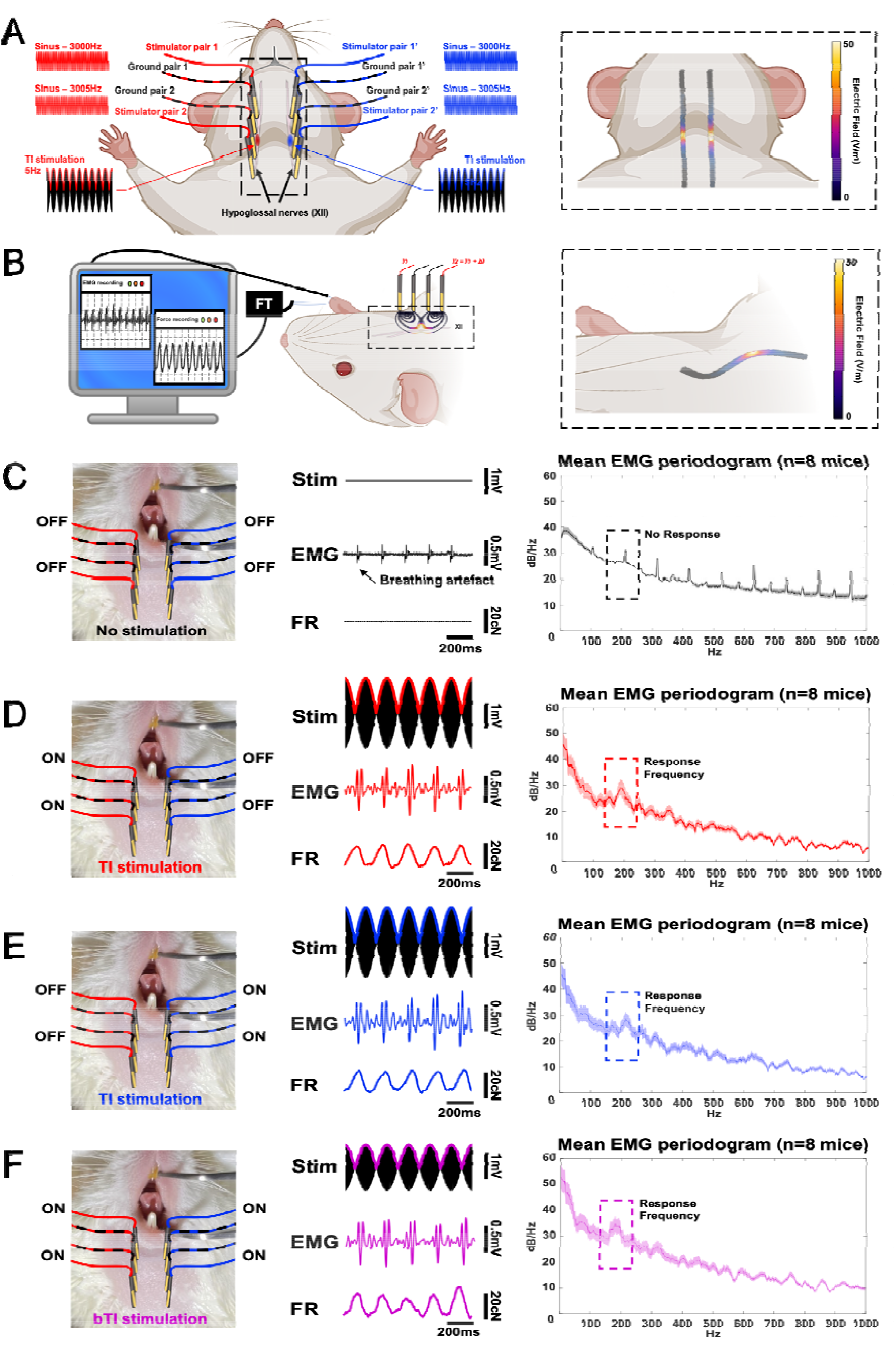
Bilateral Temporal Interfering fields for hypoglossal nerves stimulation. Bilateral TI focuses TI stimulation on both hypoglossal nerves to create a behavioral output with lower exposure strengths than regular TI (**A**). The direct output of hypoglossal nerve stimulation is tongue movement, which increases airflow and avoids sleep apnea. In our protocol, we combine tongue movement recording^7,11^ and electromyography (EMG)^10,20^ to demonstrate the value of non-invasive bilateral TI as a means for replacing implantable devices (**B**). bTI and TI simulations were simulated in Sim4Life to optimize electrode placement along the nerves and minimize unwanted stimulation in the neighboring areas. Temporal interference using a 5Hz offset efficiently stimulates hypoglossal nerves in mice. However, when stimulating one side only, the stimulation amplitude needed to elicit tongue movement and an EMG response (**D and E**) must be increased and as a result, creates unwanted stimulation of nontarget muscles. When stimulated with our bTI protocol, the stimulation amplitude of each side can be reduced, leading to selective stimulation of both hypoglossal nerves without any unwanted stimulation (**F**). The depicted EMG and FR are single-mouse examples.

With all stimulation protocols, we could induce tongue movements and record Compound Muscle Action Potentials (CMAP) in the tongue muscle. As expected, tongue movement depended on the stimulation frequency. Figure 2 shows a force recording of the tongue muscle during a 5Hz TI stimulation. To facilitate the comparison of evoked activities across the different stimulation conditions, they were normalized to those associated with a tongue movement force of about 20cN. This way, we were able to investigate differences in the stimulation amplitude needed to evoke tongue movements of the same magnitude throughout our stimulation protocols.

Comparing left and right TI, no significant differences were found in the stimulation amplitude required to elicit a 20cN tongue movement (stimulation amplitude = 2.7 ± 0.6 mA per electrode pair with no significant difference between right and left unilateral TI; p-value_Wilcoxon test_ = 0.88). However, when looking at the stimulation amplitude needed for the bTI protocol to induce a 20cN tongue movement, the required stimulation amplitude goes down to 1.7 ± 0.4 mA per pair – an amplitude significantly lower than that of the unilateral TI groups (p-value_Wilcoxon test_ = 0.01). As expected, these behavioral and electrophysiological activities were evoked only when stimulating with TI and with an offset between *f1* and *f2*. When stimulated with two identical high frequencies (*f1* = *f2* = 3000Hz), no low-frequency amplitude modulation is generated, and we could not evoke any tongue movement or CMAP signature from the tongue muscles (**Figure 3 and S1**). When stimulated with two identical low frequencies (*f1* = *f2* = 5Hz), pain sensation and involuntary muscle twitching caused by strong stimulation of skin and superficial tissues prevented reaching an amplitude at which the target nerve could be stimulated. In both cases, we could therefore not induce any tongue activity **(Figure S1)**. In summary, tongue movement and CMAPs were only evoked when using TI (i.e., offset frequencies) to stimulate hypoglossal nerves. Finally, even though unilateral TI was efficient enough to elicit tongue movements and CMAPs in tongue muscle, we showed that introducing a second TI apparatus to stimulate the two symmetrical hypoglossal nerves simultaneously significantly decreases the stimulation amplitude needed to evoke the same activity (**Figure 3**) and as a result improves stimulation selectivity by reducing unwanted stimulation of non-target tissues (**Figure S2**). Moreover, unilateral stimulation of only one hypoglossal nerve can only lead to a partial and lateral protrusion of the tongue (**Figure 5**). By introducing a second TI apparatus and stimulating both hypoglossal nerves at the same time, a complete and central protrusion (**Figure 5**) is possible, increasing the airflow recovery and decreasing the necessary amplitude to achieve it^9^

**Figure 3.**
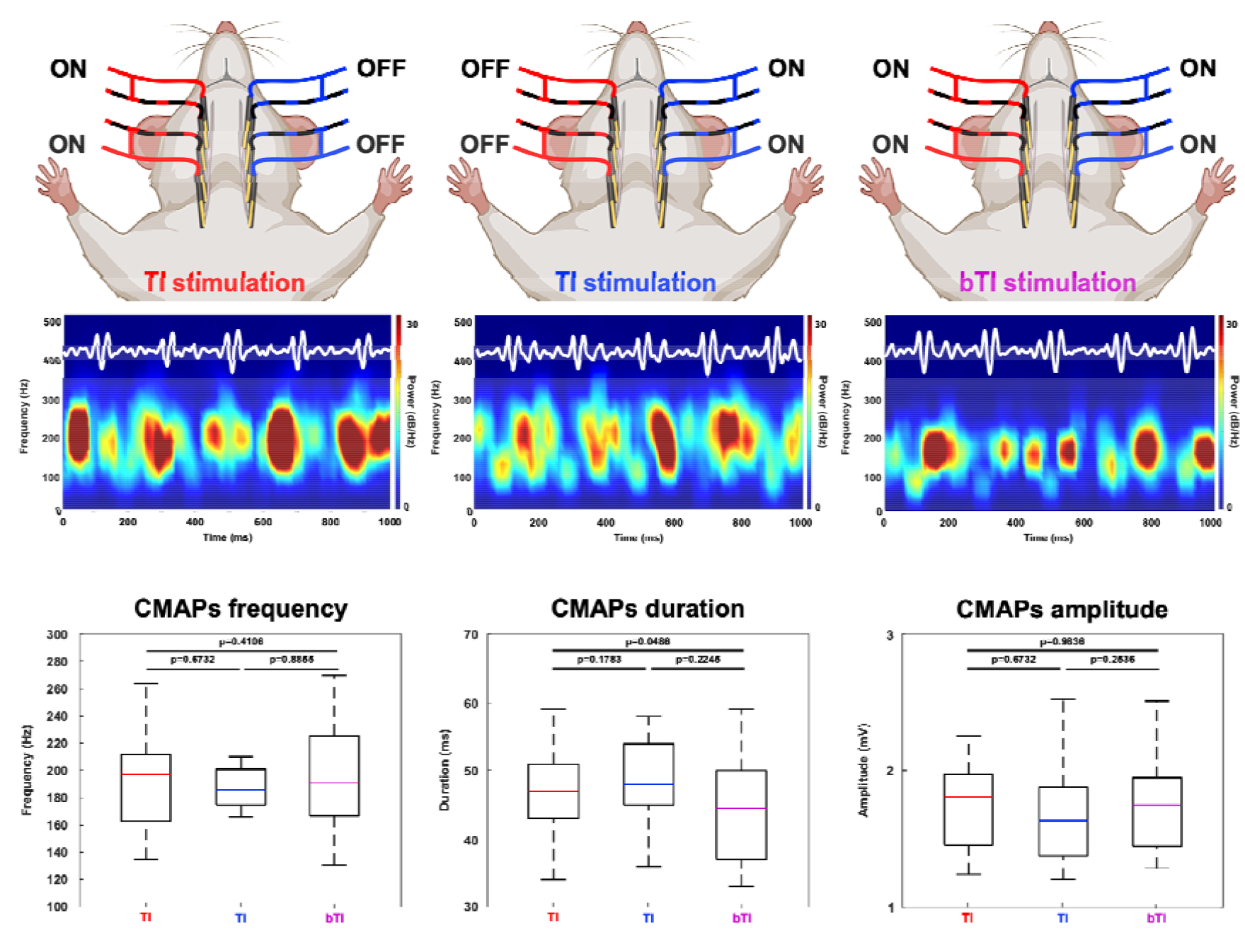
Evoked EMG response using TI and bTI. EMG response (the depicted EMG response spectra are illustrative examples from a single mouse) could be recorded in all our TI protocols. Looking at characteristics of the evoked CMAPs, we could not find any differences between unilateral TI stimulation- and the bilateral stimulation-evoked CMAPs. No significant differences between CMAP frequency components, duration, and amplitude were apparent, but the bTI required much lower stimulation amplitudes to evoke the same activity.

### Evoked Compound Muscle Action Potentials have the same characteristics in all TI protocols in mice

To be completely sure that the evoked CMAPs were identical in our three TI stimulation protocols, we collected them by stimulation groups and compared them in terms of their frequency component, duration, and amplitude. When stimulating with a 5Hz offset we can see events occurring at 5Hz that have the same global shape as tongue CMAPs following hypoglossal nerve stimulation. To make sure that the evoked CMAPs were the same for the three stimulation protocols, non-parametric Wilcoxon tests were executed to compare their frequency components, duration, and amplitude. As expected, no statistically significant differences were found between the three groups (p-values were all above 5%).

### Apnea Hypopnea Index reduction in OSA patients using bTI

During sleep studies in humans, OSA is characterized using sleep summaries and more precisely the Apnea Hypopnea Index (AHI). According to the American Academy of Sleep Medicine (AASM), apneas are defined as ≥ 90% flow reduction events lasting more than 10 seconds; hypopneas are defined as ≥ 30% flow reductions of more than 10 seconds duration with a decrease of 0_2_ saturation ≥ 3% or arousal. This way, apneas, and hypopneas can be identified, and AHI was characterized as the number of apnea events plus the number of hypopnea over the total sleep time (AHI = (# apnea + # hypopnea) / TST). Patients are then diagnosed as healthy patients (AHI<5), mild OSA patients (5<AHI<15), moderate OSA patients (15<AHI<30), or severe OSA patients (AHI>30).

In this study, 12 patients were recruited and treated overnight during an in-clinic polysomnography. Upon retrospective review, it was discovered that one patient significantly exceeded the targeted BMI range of <35 (BMI=49.8), and therefore this patient’s data were excluded from analysis, despite completing the full treatment protocol. Two other patients exceeded the BMI enrollment criteria of BMI =< 32 (33 and 33.7), but were included in the analysis as other groups have shown clinical success in patients up to a BMI of 35^16^, and Inspire, an implanted hypoglossal nerve stimulator for OSA treatment, has recently asked the FDA to expand approval to patients up to a BMI of 35. The primary endpoint was % responders (AHI reduction >50% and an AHI below 20), with an expected 50% response rate. Among all patients completing the study *i*.*e* with a BMI<35kg/m^2^ and a baseline AHI<50 (7 men and 4 women), 50% of patients (4/8) responded to the stimulation therapy with a high gender-dependent effect on therapy outcome (4/4 women versus 1/7 men) although two men had an AHI decrease of 43%. Among patients that responded to bTI stimulation, their AHI decreased by 66.59% ± 3.77% (AHI=22.53 ± 6.51 when the stimulation is off, against 7.63 ± 2.84 when the stimulation is on, p-value_Wilcoxon test_ = 0.0286). Interestingly, when looking at the amplitude threshold necessary to elicit tongue movement/tension we also see a clear difference women needed less current to evoke the same behavior compared to men (4.6 ± 1.34mA for women versus 7.86 ± 1.46mA for men). Overall, bTI stimulation led to a significant decrease in AHI for a subset of OSA patients with a clear improvement in the severity of sleep apnea.

As for secondary endpoints: Sleep time at SaO2 < 90% was not significantly reduced compared to baseline according to the Wilcoxon test (baseline 2.0 ± 0.5% and 5.9 ± 2.8%). There were no device safety-related events. Questionnaires indicated that the technology was well-tolerated and would be acceptable as a product.

## DISCUSSION

We have previously demonstrated TI stimulation to be a powerful new tool in central nervous system applications^22,21,2^. However, peripheral nerve stimulation using TI is not common^6^, especially for targeting facial nerves. This limitation of TI for facial nerve stimulation is mostly due to the special arrangement of these nerves that often overlap, making a specific and non-invasive deep stimulation hard to perform. The sensitivity of our skin limits the tolerance for electrical stimulation, and the face features some of the most sensitive skin.

The bTI stimulation, which we present in this work, relies on non-invasive stimulation of symmetrical facial nerves to help overcome sensitivity limits by reducing the exposure magnitude necessary for activation. It provides a highly specific and non-invasive stimulation of facial nerves and thus permits avoidance of surgical intervention. As already shown in the central nervous system and peripheral nerve stimulation, optimization of the stimulation geometry is crucial^22,6^.

In our study, exposure characterization and electrode placement optimization were performed using EM-simulations with Sim4Life’s electroquasistatic solver. Linear, linear interleaved, and crossing configurations were compared in terms of the selectivity and magnitude of TI HN exposure (**Figures 4 and 5**). Electrode arrangement for rodent experiments was based on in silico simulations and previous experiments using TI for sciatic nerve stimulation^6^, electrodes were thus placed along the hypoglossal nerves to create a TI field along it. As human inter-individual differences are much higher than in laboratory mice, we had to adapt our electrode placement to take into account small shifts in hypoglossal nerve location along all participants. In addition to in silico simulations using Sim4Life, pre-tests were realized to have a great reproducibility of nerve stimulation along humans using only one electrode placement, we ended up with an X-shaped electrode placement which increases TI field at depth and facilitate hypoglossal nerve stimulation even with an offset in its location. This way, using bTI to stimulate both hypoglossal nerves, we could elicit both behavioral (tongue movement) and electrophysiological (tongue CMAP) responses without invasively implanting a cuff electrode around nerves. Moreover, when comparing our bTI protocol with regular TENS, we showed that our bTI protocol can evoke activity when TENS cannot. This superiority of bTI over TENS supports our idea that bTI can easily be incorporated into a non-invasive treatment protocol for OSA.

**Figure 4.**
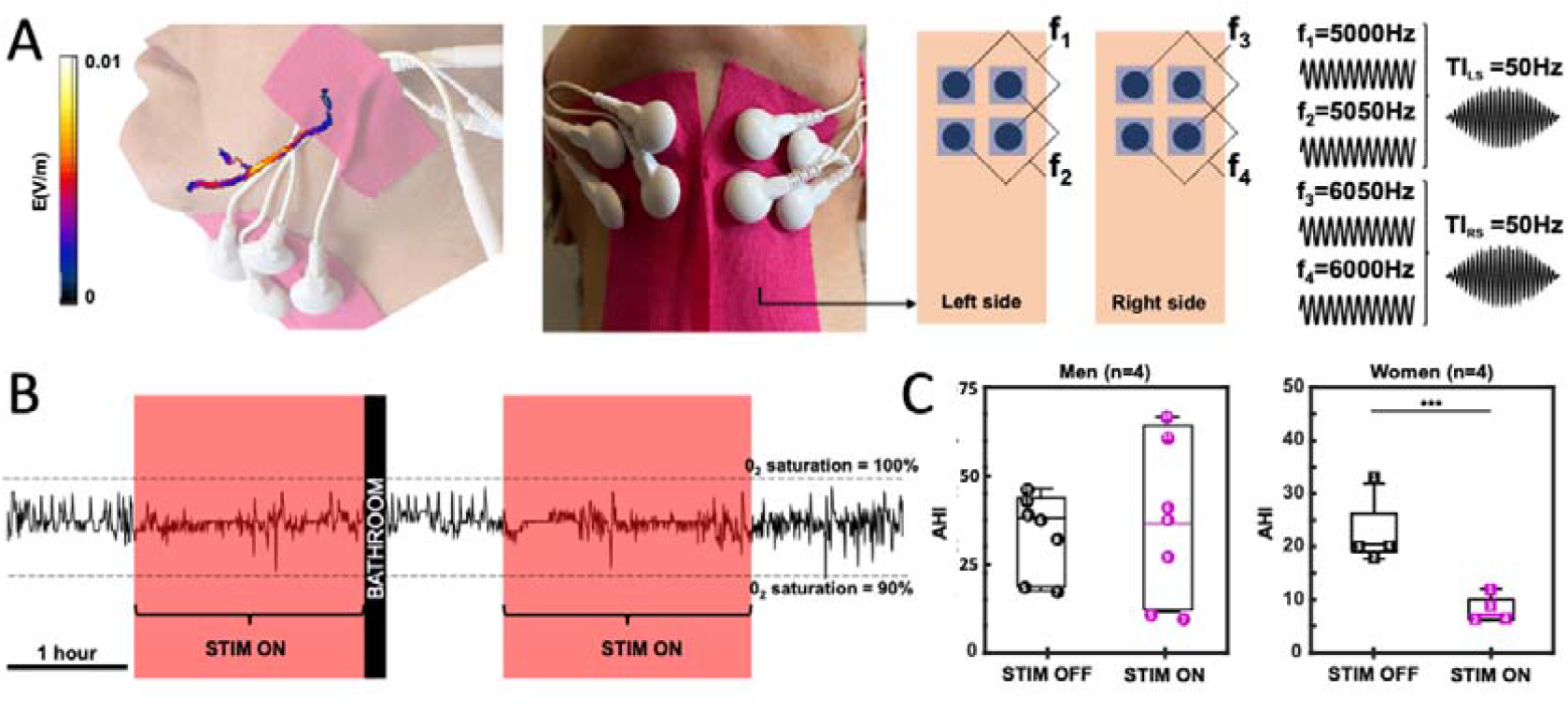
Non-invasive stimulation of both hypoglossal nerves in OSA patients reduces AHI in women. Temporal interference has been mostly applied to the Central Nervous System (CNS). As for CNS stimulation, TI can be used as a non-invasive tool to stimulate the Peripheral Nervous System (PNS) in humans. As cranial nerves exist in pairs, bilateral TI can simultaneously stimulate both cranial XII nerves focally and elicit a tongue movement or at least tension in the tongue. Simulations and experimental pre-tests allowed the identification of optimized exposure parameters and electrode placements for selective HN stimulation: 5kHz and 6kHz frequency carriers to minimalize tingling sensation on the skin when the stimulation if turned on, 50Hz TI stimulation for both HN and electrode placement around the bilateral digastric anterior belly muscles (**A**). In laboratory respiratory summaries were realized in already diagnosed OSA patients, a polysomnogram with bTI stimulation was also recorded on night 2 for each patient (night 1: baseline without bTI; night 2: recording with bTI) (**B**), and AHIs were calculated and compared to establish the severity of OSA with and without the bTI stimulation. In the present analysis, only participants with a BMI <35kg/m^2^ and a baseline AHI <50, which includes 4 women (squares 1 to 4 in boxplots) and 4 men (circles 5 to 8 in boxplots). No significant differences between stimulation ON and OFF were detected for men (p-value_Wilcoxon test_ = 0.89), whereas women seem to respond significantlyto the treatment with a clear decrease in their AHI values (p-value_Wilcoxon test_ = 0.029) (C).

**Figure 5.**
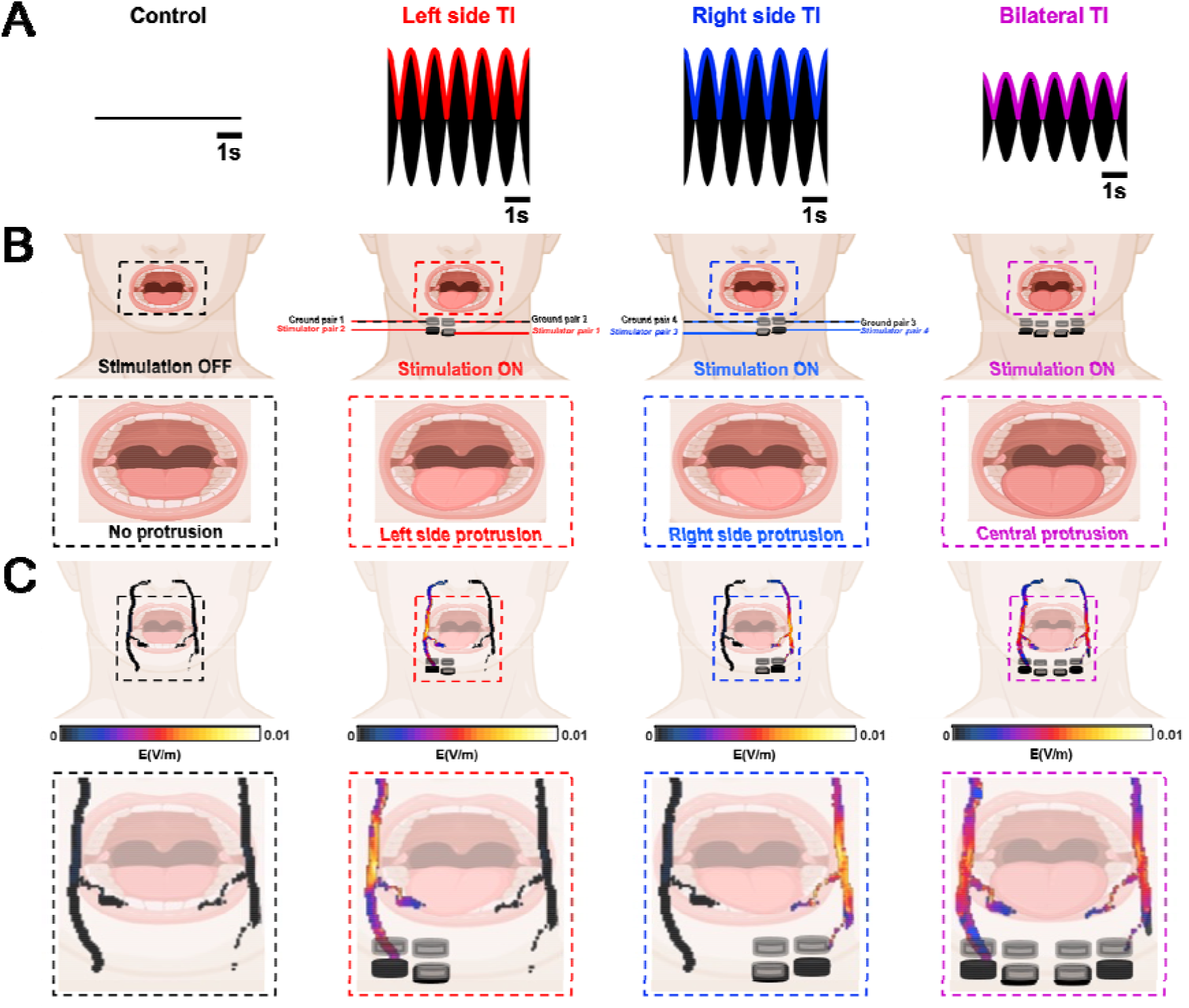
Bilateral hypoglossal nerve stimulation reduces stimulation amplitudes and induces central tongue protrusion.

Hypoglossal nerve stimulation treatment of OSA in humans is currently achieved using a cuff electrode directly placed on the hypoglossal nerves through invasive surgeries for electrode and battery implantation^19^. Using bTI for OSA patients in the clinic could avoid the use of an invasive device and provide stimulation that is as or more efficient than classical *in situ* stimulation. Instead of stimulating only a small part of the hypoglossal nerves, clinicians would be able to stimulate along the nerves and optimize nerve depolarization. Furthermore, unilateral hypoglossal nerve stimulation creates a lateral protrusion of the tongue. Using bilateral stimulation of both hypoglossal nerves allows reduction of each stimulator’s amplitude, but also induces a central protrusion of the tongue which is known to optimize airflow (**Figure 5**.).

Future bTI protocols could be performed in combination with novel electrode technologies, such as organic electronics, to make them safer and more suitable for long-term experiments^5,6,22^.

While primary clinical endpoints regarding responder rates were not achieved for the total subject population, retrospective analysis of the data revealed a strong gender bias toward females in the responder group (**Table 2**). This result is consistent with observations in long-term studies of implantable hypoglossal nerve stimulators that being female is a strong predictor of surgical success^17,28^. Although gender differences have been reported, as in the present study, no explanation has been yet exposed. To date, no anatomical differences (besides high BMI that might impact nerve stimulation, have been shown between males and females for hypoglossal nerve stimulation. In the present study, it has been shown that man requires more amplitude of stimulation to elicit some tonus in the tongue (or a tongue movement), with little flexibility left for amplitude adjustment during the night. This difference in stimulation amplitude could explain why electrical stimulation leads to less beneficial effects for men. The sample size in this study was small, and further studies with larger sample sizes are necessary to further elucidate these gender differences. In addition, this study was undertaken on a single night. Generally, therapies involve a long-term adjustment and ramp-up period. Further studies of long-term chronic stimulation are needed to determine, whether patients will undergo physiological changes to their tongues that may improve therapy after chronic stimulation, whether desensitization allows increases in energy supply, creating a stronger effect.

**Table 2.** Subgroup analysis. *** shows a significant decrease in AHI using a non-parametric Wilcoxon test.

| Subgroup | Sex<br>(♂/♀) | Age | BMI | Baseline AHI | Treated AHI | % reduction |
| --- | --- | --- | --- | --- | --- | --- |
| All | 5/7 | 48.1 ± 10.7 | 30.6 ± 6.8 | 30.1 ± 11.3 | 29 ± 24.2 | 16.3 ± 50.9 |
| Female | 5/0 | 51.2 ± 8.6 | 32.9 ± 10.5 | 24.5 ± 7.2 | 17 ± 21 | 39.9 ± 59.9 |
| Female, BMI<br>< 35 | 4/0 | 50.5 ± 9.8 | 28.7 ± 5.2 | 22.5 ± 6.5 | 7.6 ± 2.8 | 66.6 ± 3.8<br>(***) |
| Male | 0/7 | 47.9 ± 12.1 | 29.4 ± 2.4 | 34.1 ± 12.4 | 37.7 ± 23.9 | -0.5 ± 39.5 |
| BMI < 32 | 2/6 | 47.1 ± 12.9 | 27.7 ± 2.8 | 30.5 ± 11.8 | 27.2 ± 22.6 | 21.2 ± 42.5 |
| BMI < 35 | 4/7 | 47.5 ± 8 | 29.1 ± 2.6 | 29.9 ± 10.3 | 26.7 ± 20 | 23.9 ± 39.5 |

## CONCLUSION

Taken together, the data in this study support the mechanistic and therapeutic framework for the use of noninvasive bioelectronic devices to target neural circuits previously mapped to control tongue movement. Bilateral TI warrants further study for the treatment of OSA and other disorders, including systemic inflammation, and may be of particular relevance in patients that do not tolerate implantable stimulators or for whom they are otherwise contraindicated or unavailable.

## LIST OF ABBREVIATIONS

AASM: American Academy of Sleep Medicine
AHI: Apnea Hypopnea Index
bTI: Bilateral TI
BMI: Body Mass Index
CNS: Central Nervous System
CFR: Code of Federal Regulations
CMAP: Compound Muscle Action Potentials
EMG: Electromyography
FEM: Finite element modeling
FR: Force recording
HNS: hypoglossal nerve stimulation
ICH GCP: International Conference on Harmonisation Good Clinical Practice
OSA: Obstructive sleep apnea
PEC: Perfect electrical conductors
PNS: Peripheral Nervous System
PSG: Polysomnogram
TI: Temporal interference
TST: Total sleep time
TENS: Transcutaneous Electrical Nerve Stimulation

## DECLARATIONS

### Ethics approval and consent to participate

Mice experiments were approved by the European Council Directive EU2010/63 and French Ethics committee (comité d’éthique en experimentation animale n°70 Williamson, n. APAFIS 20359 - 2019041816357133). Regarding human clinical trial, it was realized with the agreement of the International Conference on Harmonisation Good Clinical Practice (ICH GCP), applicable United States Code of Federal Regulations (CFR) and followed the approved BRANY IRB File # 22-02-636-1279. All participants were given an informed consent before the trial explained all procedures and expected outcome of treatment or potential adverse effects (no adverse effects were seen during this trial).

### Consent for Publication

Not applicable

### Availability of data and materials

The datasets used and/or analysed during the current study are available from the corresponding author on reasonable request

### Author contributions

A.W. conceived the project. F.M., M.S.E., J.T. and L.S. performed experiments and F.M. analyzed neural data. I.N., J.T., E.D.G., E.N., and A.M.C. did the finite element modeling. F.M. and A.W. wrote the paper with input from the other authors including I.N., M.J.D., E.A., B.B., E.D.G., W.M.H, E.N., and L.S..

## Acknowledgments

Not applicable

## FIGURES

**Figure S1.**
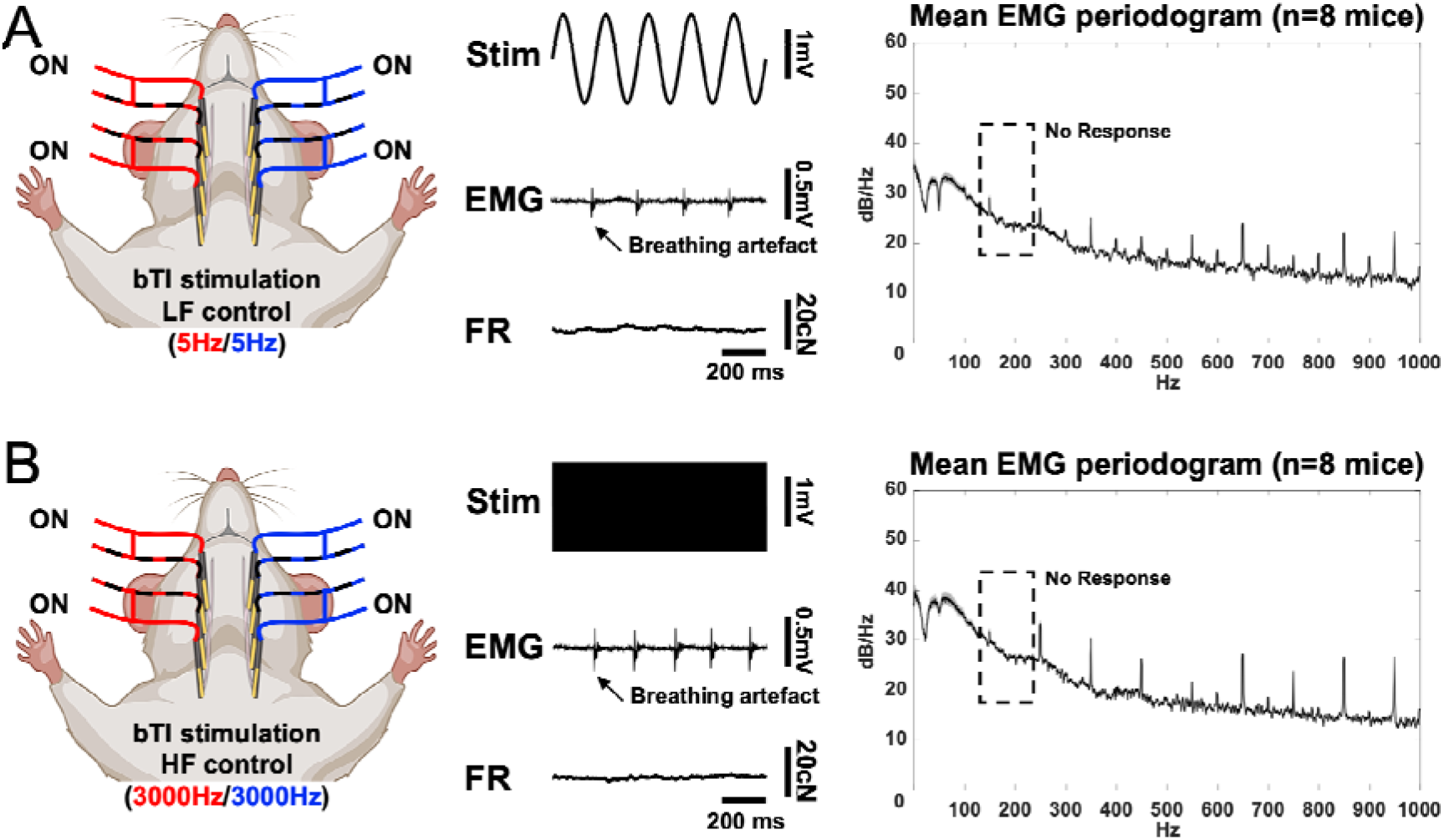
Positive high and low frequency controls bTI.

**Figure S2.**
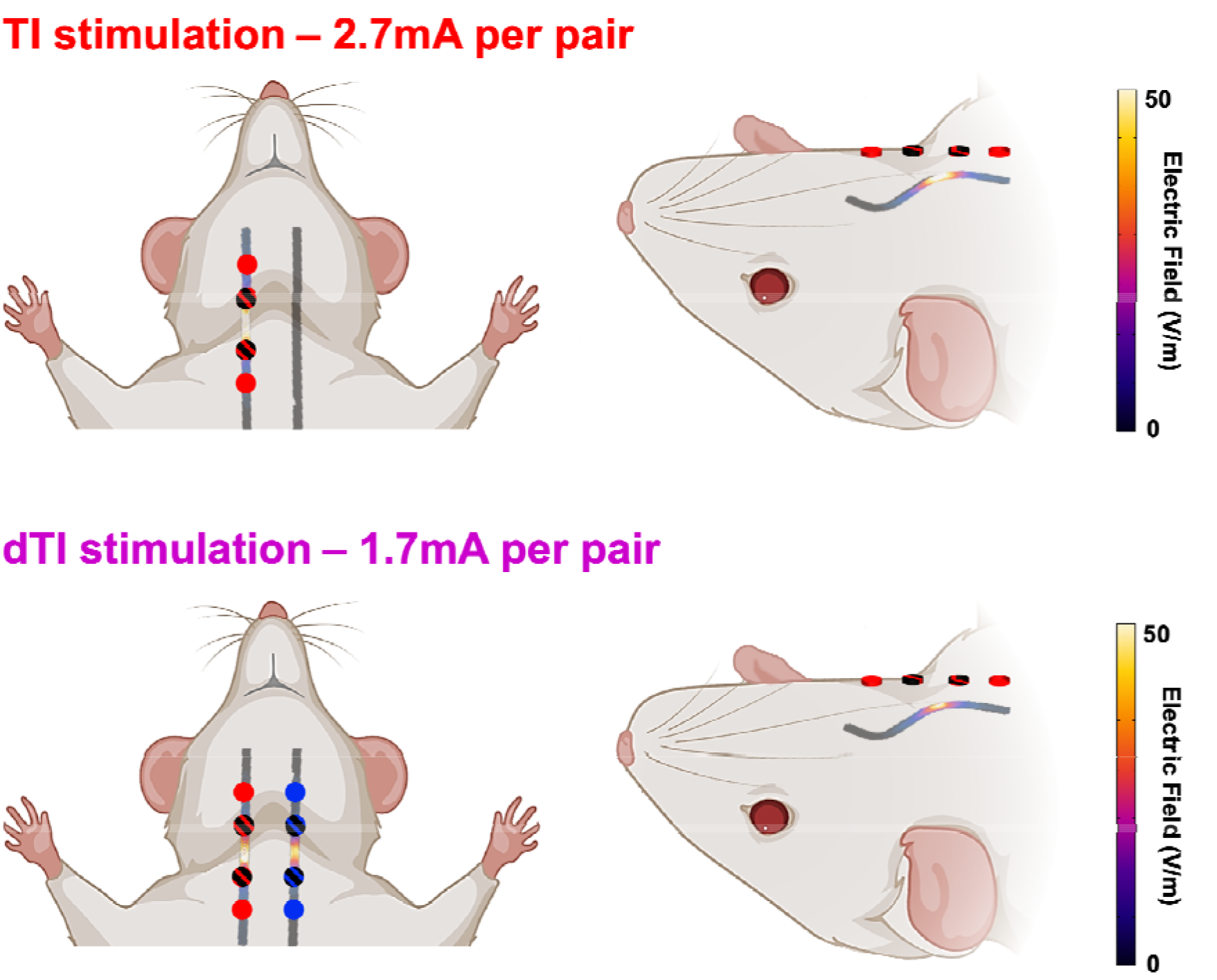
Finite element simulation comparison of bTI unilateral TI exposures.

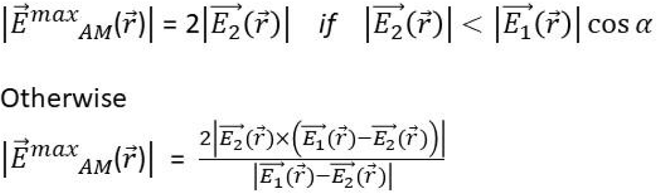

**Equation 1. Amplitude modulation formula.**

## REFERENCES

1. Abbasi A, Meghrajani V, Sharma S, Kamholz S. A comprehensive review of obstructive sleep apnea. :13.

2. Acerbo E, Jegou A, Luff C, Dzialecka P, Botzanowski B, Missey F, et al. Focal non-invasive deep-brain stimulation with temporal interference for the suppression of epileptic biomarkers. Front Neurosci. 17 août 2022;16:945221.

3. Aurora RN, Casey KR, Kristo D, Auerbach S, Bista SR, Chowdhuri S, et al. Practice Parameters for the Surgical Modifications of the Upper Airway for Obstructive Sleep Apnea in Adults. Sleep. oct 2010;33(10):1408–13.

4. Baptista PM, Costantino A, Moffa A, Rinaldi V, Casale M. Hypoglossal Nerve Stimulation in the Treatment of Obstructive Sleep Apnea: Patient Selection and New Perspectives. NSS. févr 2020;Volume 12:151–9.

5. Birmingham K, Gradinaru V, Anikeeva P, Grill WM, Pikov V, McLaughlin B, et al. Bioelectronic medicines: a research roadmap. Nat Rev Drug Discov. juin 2014;13(6):399–400.

6. Botzanowski B, Donahue MJ, Ejneby MS, Gallina AL, Ngom I, Missey F, Acerbo E, Byun D, Carron R, Cassarà AM, Neufeld E, Jirsa V, Olofsson PS, Głowacki ED, Williamson A. Noninvasive Stimulation of Peripheral Nerves using Temporally-Interfering Electrical Fields. Adv Healthc Mater. 2022;11:2200075. doi: 10.1002/adhm.202200075.

7. Connor NP, Ota F, Nagai H, Russell JA, Leverson G. Differences in Age-Related Alterations in Muscle Contraction Properties in Rat Tongue and Hindlimb. J Speech Lang Hear Res. août 2008;51(4):818–27.

8. Donovan LM, Boeder S, Malhotra A, Patel SR. New developments in the use of positive airway pressure for obstructive sleep apnea. Journal of Thoracic Disease. 2015;7(8):20.

9. Eastwood PR, Barnes M, MacKay SG, Wheatley JR, Hillman DR, Nguyên XL, et al. Bilateral hypoglossal nerve stimulation for treatment of adult obstructive sleep apnoea. Eur Respir J. janv 2020;55(1):1901320.

10. Fleury Curado T, Fishbein K, Pho H, Brennick M, Dergacheva O, Sennes LU, et al. Chemogenetic stimulation of the hypoglossal neurons improves upper airway patency. Sci Rep. juin 2017;7(1):44392.

11. Gilliam EE, Goldberg SJ. Contractile properties of the tongue muscles: effects of hypoglossal nerve and extracellular motoneuron stimulation in rat. Journal of Neurophysiology. 1 août 1995;74(2):547–55.

12. Grossman N, Bono D, Dedic N, Kodandaramaiah SB, Rudenko A, Suk H-J, Cassara AM, Neufeld E, Kuster N, Tsai L-H, Pascual-Leone A, Boyden ES. Noninvasive Deep Brain Stimulation via Temporally Interfering Electric Fields. Cell. 2017;169(6):1029–1041.e16. doi: 10.1016/j.cell.2017.05.024.

13. Günter C, Delbeke J, Ortiz-Catalan M. Safety of long-term electrical peripheral nerve stimulation: review of the state of the art. J NeuroEngineering Rehabil. Déc 2019;16(1):13.

14. Hasgall PA, Neufeld E, Gosselin MC, Klingenböck A, Kuster N. IT’IS Database for thermal and electromagnetic parameters of biological tissues, Version 2.2. September 26th. 2011.

15. Heiser C, Hofauer B, Lozier L, Woodson BT, Stark T. Nerve monitoring-guided selective hypoglossal nerve stimulation in obstructive sleep apnea patients: NIM-Guided Hypoglossal Nerve Stimulation. Laryngoscope. 2016 Dec;126(12):2852–2858. doi: 10.1002/lary.26026.

16. Heiser C, Knopf A, Bas M, Gahleitner C, Hofauer B. Selective upper airway stimulation for obstructive sleep apnea: a single center clinical experience. Eur Arch Otorhinolaryngol. 2017 Aug;274(8):1727–1734. doi: 10.1007/s00405-016-4297-6.

17. Heiser C, Steffen A, Boon M, Hofauer B, Doghramji K, Maurer JT, Sommer JU, Soose R, Strollo PJ, Schwab R, Thaler E, Withrow K, Kominsky A, Larsen C, Kezirian EJ, Hsia J, Chia S, Harwick J, Strohl K, Mehra R. Post-approval upper airway stimulation predictors of treatment effectiveness in the ADHERE registry. Eur Respir J. 2019 Dec 5;53(2):1801405. doi: 10.1183/13993003.01405-2018. PMID: 30578349.

18. Jones I, Johnson MI. Transcutaneous electrical nerve stimulation. Continuing Education in Anaesthesia Critical Care & Pain. août 2009;9(4):130–5.

19. Manchanda S, Neupane P, Sigua NL. Upper Airway Stimulation/Hypoglossal Nerve Stimulator. Am J Respir Crit Care Med. 15 oct 2020;202(8):P23–4.

20. Meadows PM, Whitehead MC, Zaidi FN. Effects of targeted activation of tongue muscles on oropharyngeal patency in the rat. Journal of the Neurological Sciences. nov 2014;346(1-2):178–93.

21. Missey F, Donahue MJ, Weber P, Ngom I, Acerbo E, Botzanowski B, et al. Laser-Driven Wireless Deep Brain Stimulation using Temporal Interference and Organic Electrolytic Photocapacitors. Adv Funct Materials. 2 juin 2022;2200691.

22. Missey F, Rusina E, Acerbo E, Botzanowski B, Trébuchon A, Bartolomei F, et al. Orientation of Temporal Interference for Non-invasive Deep Brain Stimulation in Epilepsy. Front Neurosci. 7 juin 2021;15:633988.

23. Pomerantz J, Freedom T, Nickley K, Bhayani M, Gerber M, Freedman N, Viola-Saltzman M. Upper airway stimulation for obstructive sleep apnea. SLEEP. 2018;41(5):0540.

24. Rotenberg BW, Murariu D, Pang KP. Trends in CPAP adherence over twenty years of data collection: a flattened curve. J of Otolaryngol - Head & Neck Surg. déc 2016;45(1):43.

25. Schwartz AR, Smith PL, Oliven A. Electrical stimulation of the hypoglossal nerve: a potential therapy. Journal of Applied Physiology. 1 févr 2014;116(3):337–44.

26. Sluka KA, Walsh D. Transcutaneous electrical nerve stimulation: Basic science mechanisms and clinical effectiveness. The Journal of Pain. avr 2003;4(3):109–21.

27. Sturm JJ, Modik O, Suurna MV. Neurophysiological monitoring of tongue muscle activation during hypoglossal nerve stimulation. The Laryngoscope. juill 2020;130(7):1836–43.

28. Thaler ER, Schwab R, Maurer JT, Soose RJ, Larsen CG, Stevens S, Stevens D, Boon M, Huntley C, Doghramji K, Waters T, Kominsky A, Steffen A, Kezirian EJ, Hofauer B, Sommer U, Withrow K, Strohl K, Heiser C. Results of the ADHERE upper airway stimulation registry and predictors of therapy efficacy. Laryngoscope. 2020 Jun;130(6):1333–1338. doi: 10.1002/lary.28286. Epub 2019 Sep 24. PMID: 31550076.

29. Young T, Peppard PE, Gottlieb DJ. Epidemiology of Obstructive Sleep Apnea: A Population Health Perspective. Am J Respir Crit Care Med. mai 2002;165(9):1217–39.

30. Zhu Z, Hofauer B, Wirth M, Heiser C. Long-term changes of stimulation intensities in hypoglossal nerve stimulation. Journal of Clinical Sleep Medicine. 15 oct 2020;16(10):1775–80.

